# MicroWorldOmics: A “One step for All” desktop application for comprehensive analysis in microbiome and virome and exploring “Dark Matter” from MicroWorld

**DOI:** 10.1101/2024.06.24.600528

**Authors:** Runze Li, Wei Dong, Mingjie Wang, Xiaoqing Hu, Jiao Wan, Zhuang Yang

## Abstract

A large amount of high-throughput sequencing data generated in ecology, medicine, and pharmacy has increasingly raised the complexity of data analysis and interpretation. However, the microbiome and virome field still lacks a convenient and programming-free desktop application for the comprehensive analysis in microbiome and virome data, particularly in virome analysis and ‘Dark Matter’ exploration. Therefore, a plugin development mode desktop service, MicroWorldOmics, is proposed with a convenient and one-stop pipeline for life sciences and biomedical fields. Plugin development allows users to analyze data in parallel and interactively, while these tasks usually require advanced bioinformatician to achieve. MicroWorldOmics is a software widely used in microbiome and virome, containing 90 sub-applications, including four main functions: epidemiological analysis, metagenomic/amplicons and virome comprehensive analysis, and exploring ‘Dark matter’. More than 80 Python modules and 600 R packages are invoked in all aspects of bioinformatics analysis, statistical analysis, deep learning, and visualization. At the same time, the software supports multiple input and output results, such as GFF3, FASTA, CSV, PNG, JPG, JSON, and TXT, etc. Besides, to improve the efficiency for users, MicroWorldOmics supports being deployed in three mainstream systems: Windows, Linux, and Mac and also sets the demo/example data for users to test the benchmark easily. In summary, the development of MicroWorldOmics greatly facilitates the process of analyzing microbiome and virome data for researchers who have no programming foundation and are devoted to life science and biomedicine data analysis.

## Introduction

Over the past two decades, the development of high throughput technology such as metagenomic sequences provided a solution to biological problems. Current mainstream sequencing technologies, including ‘Illumina sequencing’, ‘ThermoFisher’s Ion Torrent’, ‘PacBio SMRT sequencing’ and ‘Oxford Nanopore Technology (ONT)’ [1] have been widely applied in ecology, medicine, and Environics. The advances in high-throughput sequencing technologies have provided a new perspective to microbial world analysis at a global scale, for instance, based on deep metagenomic sequencing, the evolutionary dynamics of multi-ethnic reproductive tract microbiome have been revealed [2], Genome-resolved metagenomics of environmental samples allows the discovery of new biosynthetic gene clusters [3]. The construction of the Unified Ruminant Phage Catalogue (URPC) [4] and Chinese Human Gut Virome (CHGV) [5] is helpful in understanding the process of gene exchange between phage and host bacteria and also provides a high-precision data set for the confirmation of phage-host relationship. With the ‘microworld’ field being paid more attention and more related data generated, methods and tools for integrating data analysis need to become more efficient and convenient. Especially in the face of complex data types (FASTQ, FASTA, matrix, etc), visualization methods, and differences in analysis methods, how to create a diversified, personalized, and convenient analysis platform or software has become one of the difficulties in the development of biological information tools.

Bioinformatics is a frontier interdisciplinary discipline in the field of life science in the 21st century, which consists of mathematics, computer science, and life science, aiming at exploring life activities at all levels of transmission of genetic information from DNA to proteins. Traditional methods of qualitative data analysis are not conducive to analyzing large amounts of data generated by high throughput sequence, high dimensions of the data, difficulty of programming, and cross-platform software have become huge obstacles for scientists to explore the mysteries of biology from massive data. Benefiting from the large amount of biological data generated, more and more excellent analysis platforms have been developed. Development environment from the analysis platform indicated that application types were divided into three categories: (1). Based on community development: Bioconda [6] (https://bioconda.github.io/), Biopython [7] (https://biopython.org), Bioperl [8] (https://bioperl.org/), Biojava [9] (https://biojava.org/), Bioconductor [10] (https://www.bioconductor.org/)and CRAN [11] (https://cran.r-project.org/). Community-based platforms like those offer a large number of bioinformatics analysis packages while requiring some programming ability from the user. (2). Based on Web-service: Web analysis platforms such as Galaxy [12] (https://usegalaxy.org/), Hiplot [13] (https://hiplot.cn/) and TOmicsVis [14] (https://shiny.hiplot.cn/tomicsvis-shiny/)provide users with convenient online analysis services, accompanied by high consumption of computing resources, requiring long-term maintenance services. (3). Desktop application development. The software represented by TBtools [15], OmicsSuite [16], eGPS [17], and PhyloSuite [18] provide localized, convenient, and efficient biometric information analysis tools, which do not require costly server maintenance, and protect the privacy of user data. In summary, existing technologies and approaches for bioinformatics analysis provide an excellent platform for the development of new applications that would enable to analyze a wide range of biological data, however, it lacks personalized analysis service in certain field.

According to the analysis requirements in the field, facing for microbiome and virome desktop suite named “MicroWorldOmics” was developed. In this application, friendly microbiome and virome multi-omics analysis and visualization were provided, which included 90 sub-applications (self-development and public applications), conducting epidemic analysis of pathogenic bacteria, phage taxonomy assignment, core microbial population identification, microbial interaction network construction analysis, etc (public applications). Meanwhile, self-developed algorithms phage “Dark matters” mining, microbial core genome construction, pathogen serotype identification, and other functions (self-development) have also been added to “MicroWorldOmics". “MicroWorldOmics” provides facing for microbiome and virome one-stop workflow which can be easily applied to fields of basic medicine, veterinary medicine, environmental science, and ecology. At present, the development details and download address of “MicroWorldOmics” are published at https://github.com/hzaurzli/MicroWorldOmics.

## Materials and methods

### Concise and easy-to-use desktop interfaces and public plugins

The classic UI development framework, QTdesigner (5.11.1), using PYQT5 syntax and an in-house UI component library to provide a reactive, was integrated into constructing the desktop interfaces. A primary advantage of the designed MicroWorldOmics was providing a user-friendly and efficient desktop application through a graphical user interface. Therefore, this application shared a horizontal layout of the web page, the top panel and left panel showed the thumbnail of the plugins and allowed users to quickly link to the corresponding plugins. The application logo, development information, and logs were exhibited in the center panel and right panel. Consequently, there is the possibility to perform efficient multitasking parallel on multiple working pages because of plugin development.

Multiple forms of data import were supported as the core interface of MicroWorldOmics, including GFF3, FASTA, CSV, PNG, JPG, JSON and TXT, etc, which was developed based on the PYQT5, using the QtWidgets framework. Meanwhile, this function has been wrapped into each standalone plugin to facilitate easier integration in MicroWorldOmics. In addition, Shiny apps were also embedded in MicroWorldOmics as one of the featured modules, which were stimulated by the linkage of PYQT5 and Rscript to launch the execution of Shiny apps. Based on the rendering elements of hypertext files by PYQT5, the conveniently HTML-based plugins on MicroWorldOmics embodied the powerful visualization performance and obtained high-quality output in the browser window. MicroWorldOmics has developed a stable desktop application based on PYQT5 and plugins for “Micro-World” multi-omics data analysis based on the QtWidgets framework. Combining Python’s excellent performance in deep learning, powerful data statistics, and visualization capabilities in R, a total of 90 sub-applications and more than 600 R packages and Python modules were integrated into MicroWorldOmics and provided convenient installation package, users do not need secondary configuration after installation. For all plugins in MicroWorldOmics, all applications were collected in PyPI (https://pypi.org/), Bioconda [6] (https://bioconda.github.io/), Biopython [7], Bioconductor [10] (https://www.bioconductor.org/), CRAN [11] (https://cran.r-project.org/) and GitHub (https://github.com).

### Cross-platform plugins construction

The cross-platform plugin invocation was controlled based on the Python (V 3.6) programming language (https://www.python.org), which conferred great advantages in helping users to quickly and simultaneously analyze multiple data sets in different environments. Besides, for Windows users, two modes were performed to execute Linux software, the mode of remote server transfer was adopted in MicroWorldOmics and the initial files could be uploaded to a remote server for analysis, such as Genomad [19] (https://github.com/apcamargo/genomad/) and Prokka [20] (https://github.com/tseemann/prokka). Another mode was adopted was that relevant Linux software were compiled by Cygwin (https://cygwin.com/), such as FastANI [21] (https://github.com/ParBLiSS/FastANI), CDhit [22] (https://github.com/weizhongli/cdhit) and ARAGORN [23] (https://github.com/TheSEED/aragorn). For Mac or Linux users, all software could be configured by Conda (https://www.anaconda.com/).

### Self-develop algorithm tasks

In this study, seven plugins were developed independently:

#### a. Multi-Locus Sequence Typing (MLST)

The multi-locus sequence types were determined by extracting the allele types of housekeeping genes from different isolates’ genomic sequences, all housekeeping genes in collected isolates were shown on the website PubMLST [24] (https://pubmlst.org/). BlastN V 2.15 [25] was used for detecting targeted housekeeping genes on the genome-wide scale with 100% percent and evalue < 10-5. Then the number of each housekeeping gene was counted and compared with the existent MLST available at PubMLST to obtain the sequence type of each sequenced isolate. Otherwise, a new MLST was defined.

#### b. Serotype analysis

Major antigenic variability upon the cell surface, lipopolysaccharide (LPS) O antigen confers, was used for bacteria serotyping. In our algorithm, two modes were designed: (1). Normal mode, in this mode, the different isolates’ sequences were aligned to the corresponding serotype sequences by BlastN V 2.15 [25] (evalue < 10-5), serotypes of each isolate was determined according to the comparison results. (2). Precise mode, in some special cases, such as S. suis, serotype 2 and serotype 1/2, serotype 1 and serotype 14 only differ in 161th amino acid of the cpk protein sequence. For the above situation, profiles (in the demo dataset) can be uploaded by users and note the different positions of amino acids. The mode will use the sliding window and global alignment algorithm to locate and distinguish different amino acid sites.

#### c. Genes identification

Special genetic elements’ identification is one of the important components in the pathogenic field, and a plugin for identifying genetic elements based on this requirement was designed. Based on different requirements, there are two modes designed: (1). Based on nucleic acid sequences, BlastN V 2.15 [25] (evalue < 10-5) was used to search for homologous fragments on the target contigs, users could set fragment coverage and percent in the window of the plugin according to the actual situation to get the final results. In this mode, the type of isolates query contigs and reference sequences were nucleic acids. (2). Based on protein sequences, protein fragments, which were translated from target contigs, were scanned by BlastP V 2.15 [25] (evalue < 10-5) in order to find out homologous protein fragments. fragment coverage and percent were defined by users in order to obtain satisfied results. In this mode, only protein sequences could be accepted.

#### d. Core genome construction

To identify conserved regions and functionalized elements (conjugative functionalized elements) in isolates’ genome, to achieve this, homology groups containing reference isolate CDS (coding sequence) were aligned to target isolate contigs iteratively by BlastN V 2.15 [25] (evalue < 10-5, percent is defined by users). Some homology groups were excluded from the constructed pan-genome if those homology groups did not exist in isolated contigs. In this section, identified homology groups’ positions, homology CDS, and aligned sequences were listed in the output folder.

#### e. Lysin activity evaluation

The 148,269 positive lysin sequences were collected from NCBI (https://www.ncbi.nlm.nih.gov/), which comprised from projected lysins, patents, and experimentally validated lysin sequences. The negative dataset was generated using the strategies undertaken in previous studies [26, 27]. The protein sequences with the highest annotation levels within 100-400 amino acids were downloaded from UniProt [28]. The negative dataset totally contains 3,863 lysin sequences, excluding those annotated as ‘antimicrobial, hydrolase, hemolysin, lysin, glycosidase, endolysin, holin, and bactericidal’. In order to evaluate the activity of lysins, after random positive sampling (to ensure the positive and negative samples were the same), five machine learning classifiers (see Table S1) were constructed to extract minor features in the first layer based on six features (see Table S2). Then, according to minor features extracted from the first layer, the second layer was constructed to train the evaluation model by logistic regression. Once the protein sequence was uploaded by users, the plugin would output a value between 0 and 1, the protein tends to be active if the value closes to 1.

#### f. Differential network analysis

The algorithm of differential network analysis is referred from the reference [34], here, building a differential network consists in five steps: (1). Building correlation matrix C between Operational Taxonomic units (OTU), the formula to see below:

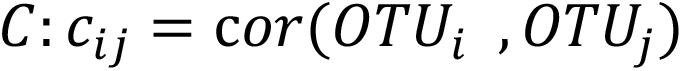

Where OUT i represents population i abundance and in this step, the Pearson correlation coefficient is implemented for subsequent analysis. (2). Computing adjacency difference matrix, different from traditional adjacency matrix in co-expression network construction, adjacency difference matrix mainly reflects the difference of microbial interaction pairs under two different conditions (correlation is used as an indicator of interaction strength).

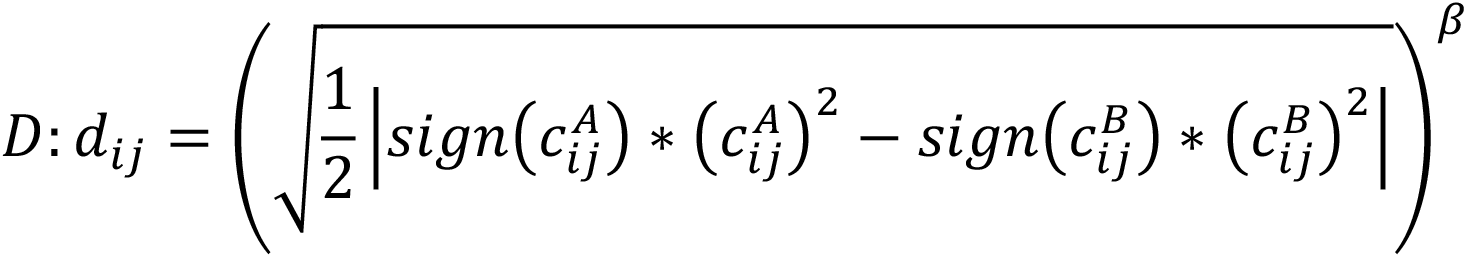

Where c^A^ represents condition A, the intensity of interaction between OTU i and OTU j, c^B^ represents condition B, the intensity of interaction between OTU i and OTU j, dij represents the difference of interaction intensity between out i and out j under two conditions (A and B), the higher the value is, the interaction intensity of OTU i and OTU j changes significantly under the two conditions. Correlation change be quantified as the difference between the square of the correlation coefficient, in order to explain the variance (r^2^) in the same correlation change and be given the same weight. β, the positive integer soft threshold to adjust for minor differences in networks, the weight of large correlation differences is emphasized by choosing a higher value, otherwise weight of lower correlation differences will be ignored. (3). Computing dissimilarity matrix T based on adjacency difference matrix D and to derive the topological overlap [35].

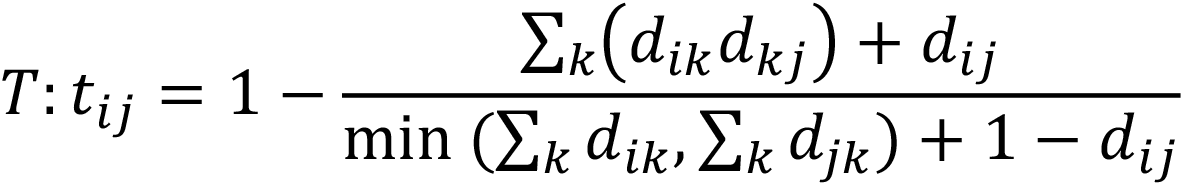

Where tij represents dissimilarity value and the tij computed here takes values between 0 and 1. The dissimilarity metrics are constructed by measuring topological overlap, which allows the identification of OTUs that share the same neighbors in the graph formed by the differential correlation network as defined by the adjacency matrix created in (2). Therefore, the lower the tij value, the higher interaction similarity between condition A and condition B, indicating that OTU i and OTU j have significant correlation changes in the same large group of OTUs. If OTU i and OTU j are part of an OTU module that is co-expressed under only one condition, then the topological overlap of OTU i and OTU j in the two conditions’ network will be different and those OTUs will be split into two modules. If OTU i and OTU j are equally correlated in both cases, then the topological overlap between OUT i and OTU j in two conditions’ network will be the same and those OTUs will exist in the same module. (4). Hierarchical clustering to split different modules. The dissimilarity matrix T is used as input to for clustering and modules are identified. The different modules are extracted from the generated tree using hierarchical clustering (dynamicTreeCut) [36]. (5). Differential network visualization.

#### g. Plaque recognition

Based on the Python module open-cv2 (https://pypi.org/project/opencv-python/), the automatic plaque recognition function was developed. More specifically, the source images were converted to several binary images, thresholding was applied, starting with minThreshold, and ending with maxThreshold in tresholdStep increments. Depending on the background color, set blobColor = 0 if the background color is dark, otherwise set blobColor = 255 and filter some plaques by setting the parameters minArea and maxArea. All parameters are set on the panel, which is provided for user settings.

## Result

### MicroWorldOmics design framework

Since the advent of metagenomic, meta-virome sequencing technology, a large number of data sets and related studies have been published. In the process of integrating, mining, and analyzing data, some obstacles for scientists are needed to overcome, such as programming skills, mathematical statistics, and visualization methods. MicroWorldOmics as an efficient application for microbiome and virome provides 90 sub-applications’ analysis services and visualization (Figure 2). All sub-applications use the PYQT5 design framework and development. PYQT5 component library was implemented for UI interface, parallel computing, web engine, and parameters passing, including some sub-libraries like “QtCore”, “QtGui”, “QtWidgets” and “Qt”. Some input and output file prompts are also inserted into the input box of each plugin for user convenience. Using the “WorkThread” module to achieve program running and UI interface separation, the function “pyqtSignal” are implemented to accept the signal that the program has finished and this design effectively avoids interface locking in computing consumable programs. Meanwhile, each plugin has a status panel that prompts for program logs and error messages.

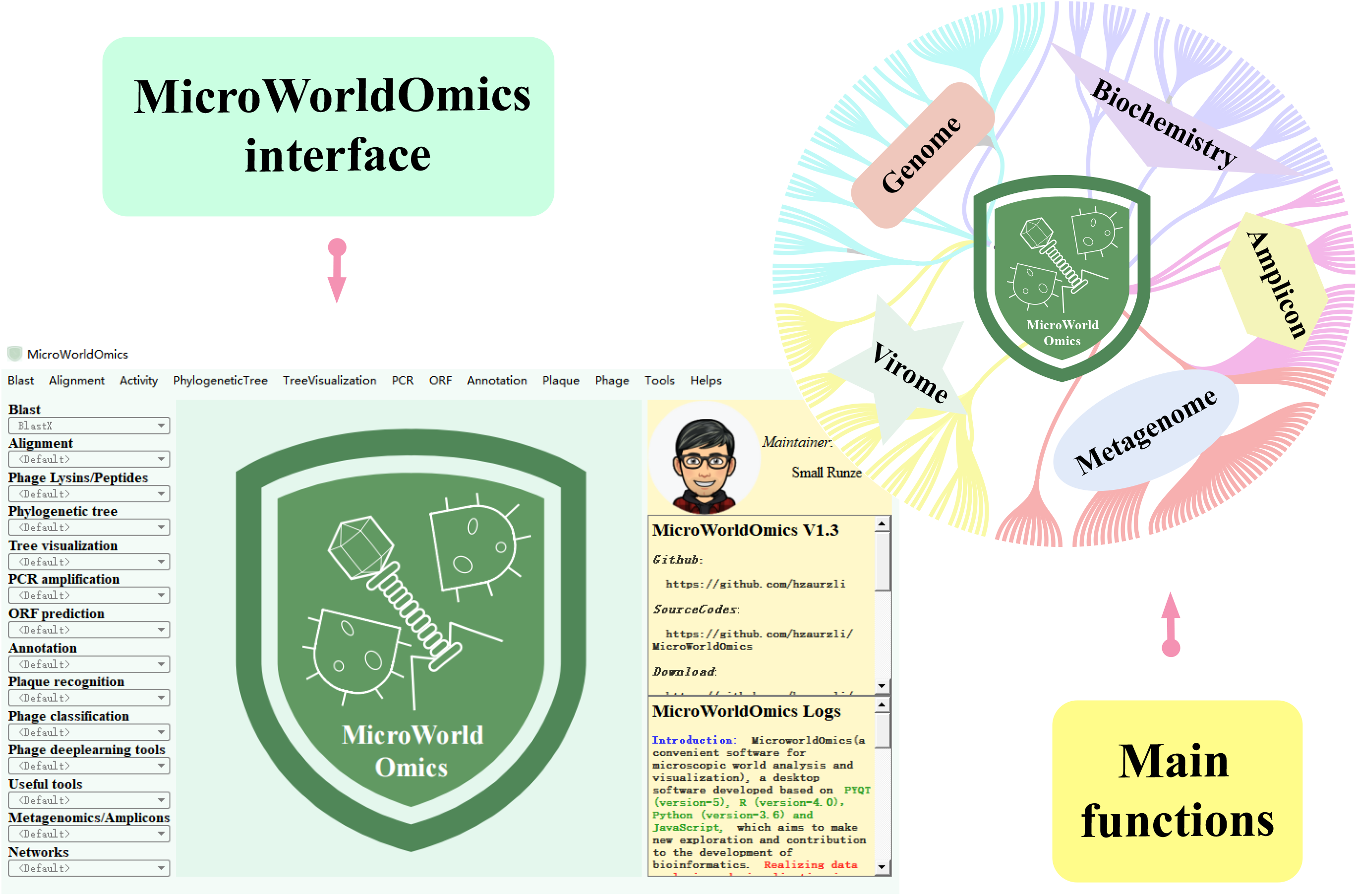
Graphical Abstract. Comprehensive analysis software in microbiome and virome implements 90 sub- applications for four main aspects, epidemiological analysis, metagenomic/amplicons, and virome comprehensive analysis and exploring‘Dark matter’. More than 80 Python modules and 600 R packages are invoked and support multiple data formats as input and output. MicroWorldOmics provides open-source download links and community services, which offer great convenience for researchers with no programming foundation.

**Figure. 2.**
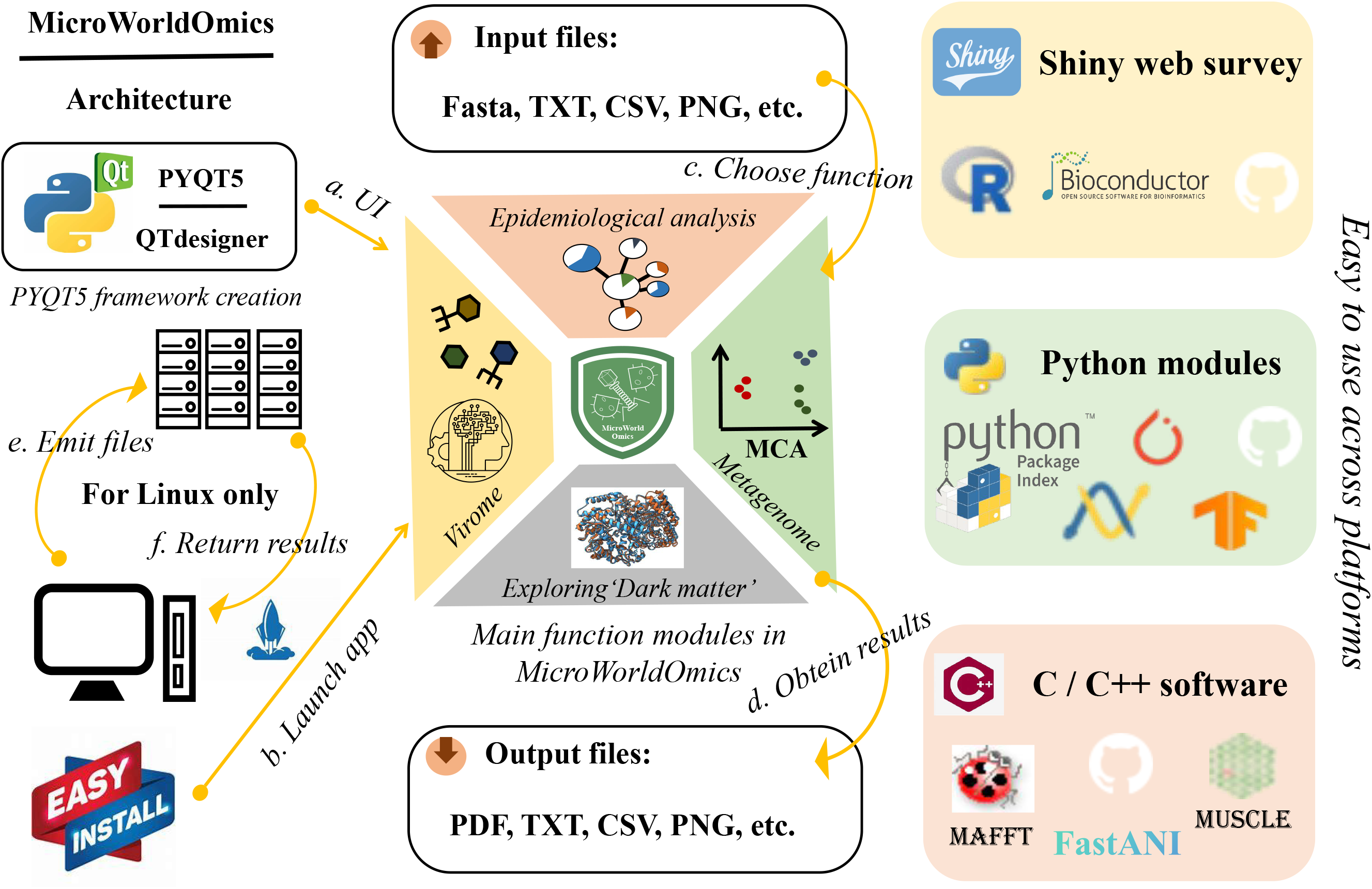
The MicroWorldOmics innovative design concept, framework architecture, and main functions. The arrows show the logical steps of execution, data set input, and result output. When the software does not support the Windows system, users can use the interface of MicroWorldOmics to upload the file to the remote server, and automatically download files to the local server after the tasks are complete.

A large number of bioinformatic modules are integrated into MicroWorldOmics, which supports multilingual software. For instance, Biopython [7] is a powerful bioinformatic analysis suite, covering the fields of sequence processing, sequence alignment, biological databases querying, structural biology, and genomics. The deep learning platforms Pytorch (1.10.1), Tensorflow (2.6.0), and Transformer (4.11.3) are used for deep learning model construction and viral data analysis. Some software written by cpp, such as Mafft [29], FastANI [21], and Muscle5 [30] can be easily modified to run on MicroWorldOmics. For some commonly used software that cannot be supported by the Windows platform, we deploy this software on a remote server, and users only need to upload data through windowing operations, then automatically download the results to the local end after running. The automated plaque counting function also brings great convenience to researchers (Figure S3), meanwhile, the structure of the chemical formula is also allowed to be drawn freely (Figure S4 A) and is provided with a variety of editable save formats (Ket format, MDL Molfile 3000, SVG, etc). Benefiting from powerful data visualization and statistical analysis capabilities, Shiny (https://github.com/rstudio/shiny) is utilized as an interface operating system by integrating lots of R packages from CRAN, Bioconductor, and Github. A total of 26 Shiny apps are integrated into MicroWorldOmics, such as R packages vegan [31] is involved in microbial diversity calculations, visualization of sequencing data requires igvShiny (https://github.com/gladkia/igvShiny), 3D protein models are constructed by r3dmol (https://github.com/swsoyee/r3dmol) (Figure S4 B), the batch effect of microbial abundance data can be easily removed using PLSDAbatch [32], R package WGCNA [33] assists in the construction of differential network. In case of windowing operation, the user can choose to start the Shiny app in MicroWorldOmics and to open the Shiny service in multiple windows. All the sample data is provided in the attachment.

In order to facilitate the needs of most users, MicroWorldOmics has developed multi-platform versions, including Windows, Linux, and Mac. For Windows, after decompressing the module package, users only need to install based on the installation file. For Linux and Mac, Users are provided with the source code and then making execution for conda to create a virtual environment. MicroWorldOmics executes excellent performance with a minimum of 8-core CPU and 8GB memory, in particular, performing deep learning calculations. At least a 4-core CPU and 4GB memory are satisfied for basic startup and some low computational resource functions. Cherry, the function with the longest single computation time, the test took approximately 3100 seconds using 20 contigs with Core (TM) i5-5200U CPU and 16GB memory.

### UI interface, design concept and analysis plugin

QTdesigner (5.11.1) was used to redesign the appearance of MicroWorldOmics to provide a convenient and comfortable environment for users. MicroWorldOmics home page shows the logo of the software in the middle, which aims to reflect the software scope: microbiome and virome. The right panel covers the development meta information, development logs, author contact information, community, etc. The top panel is a multi-level menu bar, and the left panel is a plugin menu for quick links to the corresponding plugins. The plugins menu includes several sub-plugins menus divided into 14 modules, such as “Phylogenetic tree”, “Plaque recognition”, “Phage deep learning tools”, “Metagenomics/Amplicons”, “Network”, etc. In addition, based on our extensive literature search, MicroWorldOmics is one of the comprehensive data-driven desktop services for life sciences and biomedicine, especially users can use MicroWorldOmics and its plugins to handle data analysis without operating system or software environment limitations and are also proposed to assist biology and biomedicine researchers because of convenient plugins (Table 1). The instruction for MicroWorldOmics is placed in the top panel named “help”, which can be clicked on an external web page. All introduction, source code, manual, and other information about MicroWorldOmics is reconstructed into a web page based on JavaScript Query (JQuery) and Hyper Text Markup Language (HTML), in order to establish a community to facilitate interaction between users and developers, users and users, and to better maintain MicroWorldOmics.

**Table 1.** Comparison between MicroWorldOmics and other bioinformatic platforms.

| Highlights | MicroWorldOmics | TBtools | Hiplot | ImageGP | eGPS | OmicsSuite |
| --- | --- | --- | --- | --- | --- | --- |
| Microbiome | Yes | Some | Some | No | No | Some |
| Virome | Yes | No | No | No | No | No |
| Shiny function | Yes | Yes | Yes | No | No | No |
| Deep learning | Yes | No | No | No | No | No |
| Data visualization | Yes | Yes | Yes | Yes | Yes | Yes |
| Mining ‘Dark matter’ | Yes | No | No | No | No | No |
| Pathogens analysis | Yes | No | No | No | No | No |
| Code writing | No | No | No | No | No | No |
| Self-algorithm | Yes (some) | Yes (some) | No | No | Yes (some) | No |
| Platform | Win/Linux/Mac | Win/Linux/Mac | Websites | Websites | Win/Mac/Webs | Win |

While implementing many plugins in MicroWorldOmics has enhanced the management of different functions and allowed to run programs in parallel. Now, each function plugin is provided with the corresponding sample data and each plugin has input/output and parameter types on its page, users can freely and interactively explore microworld mystery and comprehensively obtain microworld knowledge through microbiome and virome (Figure S1). When the corresponding plugin link is clicked, the layout will switch to the functional interface, most plugin functional interfaces usually contain three main sections: input/output, parameter panel, and status panel. The input/output section allows users to access example data or upload/download data through custom paths (Some plugins require users to upload files, and some plugins require users to place all files in a common folder and upload them through the folder) (Figure S2 C). The input form of the parameter panel is various, such as radio box, check box, numerical input, and image input, in order to facilitate more convenient results observation, table display functionality has also been incorporated into some plugins, and the rows and columns can be accommodated and sorted freely (Figure S2 A). Finally, the process, status, and error information of the program will be displayed in the status panel as basic tips. Other plugins launch a Shiny app via the “start” button in the interface, when the status changes to “ShinyApp has been started!!!”, then make use of the PYQT5 Web plugin to get routes and render the Shiny app by the button “Open Web". Shiny app strives to provide a concise pattern for all applications, which assists users in fully experiencing the philosophy of analysis and visualization. The input and output support multiple data formats, including TXT, CSV, PDF, JPEG, PNG, etc, users are allowed to download publishable results and images conveniently (Figure S2 B). All background, function descriptions, manuals, and citations are provided on the “help” page.

### Usage example

Owing to space limitations all the examples could not be emerged in the paper. a small number of functional plugins based on demo data sets will be presented to help readers understand the functions of MicroWorldOmics. These representative cases are divided into four major parts including epidemic analysis of pathogenic bacteria, artificial intelligence analysis in virome, metagenome-based data analysis and visualization, and network comprehensive construction.

### Case 1: Epidemiological analysis of pathogenic bacteria

In order to demonstrate the effectiveness of MicroWorldOmics in pathogenic bacteria epidemiology, The example data was collected from published research articles [37] as a case study. Multi-locus Sequence Typing (MLST), Serotype, and drug resistance analysis, the highest high-frequency research hot spot in this field, were selected to exhibit powerful usage of basic analysis tasks. After genome assembly is completed, MLST analysis was performed by the plugin “MLST”, After collecting different isolates’ housekeeping genes from a public database (PubMLST, https://pubmlst.org/), this function enables users to upload target isolate genome and extra files (housekeeping genes sequences and MLST libraries) to analyze. Subsequently, the program running in the background is called by this plugin with a tabular form as output. Meanwhile, running processes and other information can be displayed in the “status panel", after the program completion, all the results will be output to the specified path. The output results can be used as input to various drawing software, such as R (https://www.r-project.org/) and GraphPad (https://www.graphpad.com/) and the results showed that the main ST types of Streptococcus suis (S. suis) were ST1 and ST7. (Figure 3A). For some pathogenic bacteria, scientists pay more attention to their serotype, a genetic marker that exists in the blood. MicroWorldOmics also provides a plugin to support such analyses. Different from the MLST analysis plugin, serotype analysis adopts a dual-mode approach, which is divided into normal mode and precise mode. Normal mode allows users to specify different serotype sequences to do serotyping, however, in S. suis typing, serotype 1 and serotype 14, serotype 2 and serotype 1/2 differ only at position 161 of the “cpk” protein sequence, serotype 1 and serotype 2 are tryptophan, serotype 14 and serotype 1/2 are cysteine.

**Figure. 3.**
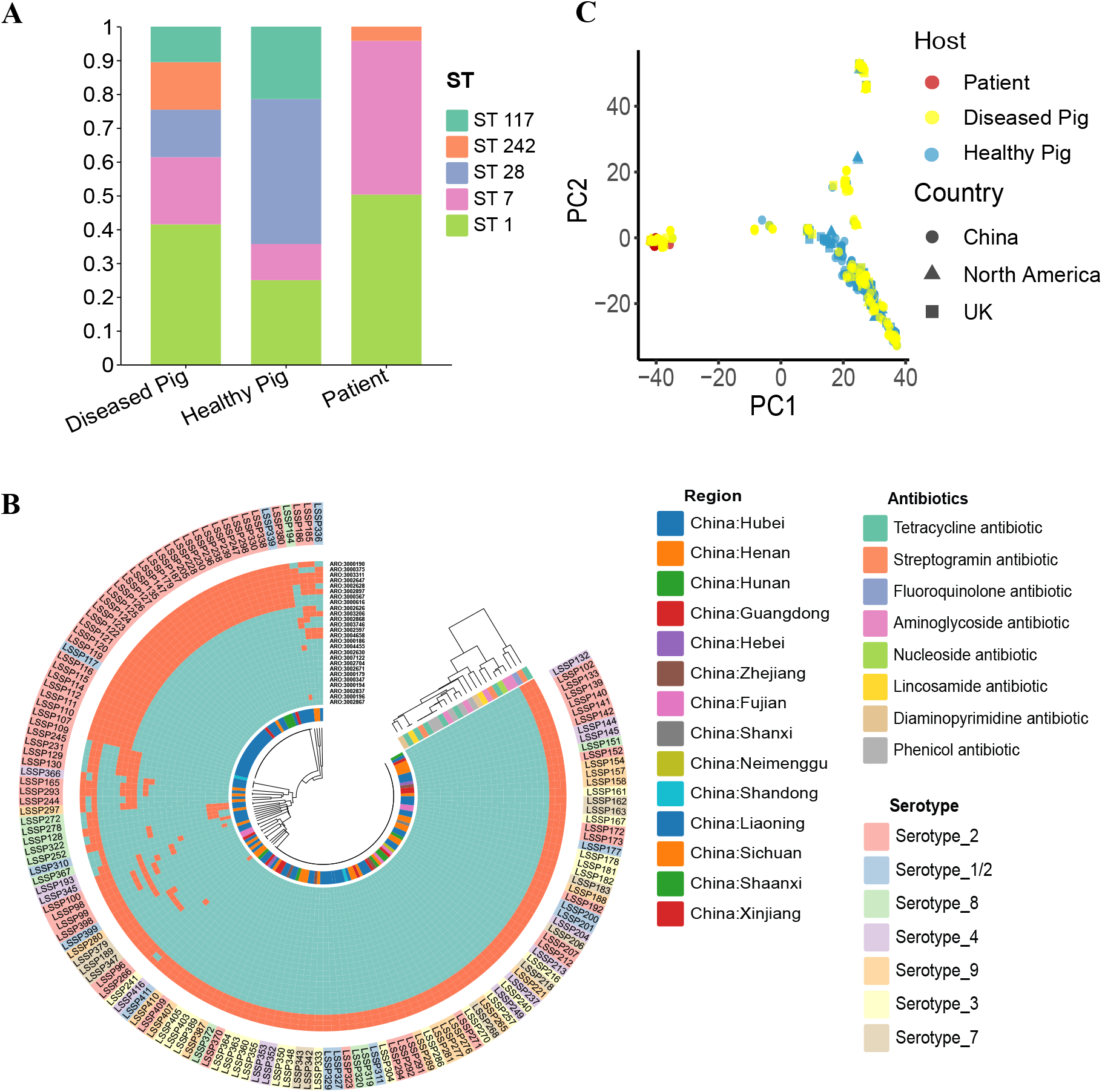
Representative use cases of pathogenic bacteria epidemiological analysis in MicroWorldOmics. (A) Graph showing the Multi-locus sequence typing (MLST) application, demo output showing an epidemic trend of ST type strains from different sources (diseased pig, healthy pig, and patients). (B) The graph shows the resistance gene carrier of demo isolates, serotype distribution, and sampling area distribution. All isolates carried fluoroquinolone antibiotic-resistance genes. (C) After obtaining all isolates’ core genomes, PCA dimensionality reduction clustering is performed using the plugin ShinyGenomePCA. S. suis from patients was clearly distinguishable from S. suis from healthy pigs.

Therefore, in precise mode, a profile table and corresponding sequence like “cpk” are specified by users to subdivide detailed serotypes with minor differences. In pathogen analysis, identification and localization of drug resistance genes are the main focus in the pipeline. Here, different treatment strategies were proposed based on the drug resistance types of different strains. Identification of drug resistance genes helps us understand the impact of new drug resistance trends and new ‘Superbugs’ discoveries. After downloading the drug resistance gene library from a public database such as CARD (https://card.mcmaster.ca/download), the “Gene Identification” plugin is used to identify different resistance genes in each isolate by controlling threshold parameters “Identification” and “Gene coverage”. Combined results of serotype and drug resistance, in 150 demo isolates, A total of 7 serotypes were identified (serotype 2, serotype 1/2, serotype 3, serotype 4, serotype 7, serotype 8, and serotype 9) and all isolates carried fluoroquinolone antibiotic resistance genes (Figure 3B).

Apart from typing study, the Shiny-based application plugin can be used to visualize large-scale isolates genome distance interactively. For example, after obtaining core elements between isolates, the sequence information is converted to a digital signal, and dimensionality is reduced by using PCA through the function “glPca()”. This plugin visualizes isolate distance using a PCA plot (Figure 3C), which may help comparison in different isolates from different isolated hosts, different regions, different genetic backgrounds, etc. After obtaining the final results, users can interactively download editable figures in the plugin.

### Case 2: Use artificial intelligence to assist virome analysis

Phages, the most abundant species of virus on Earth, the advanced analysis aims to explore and identify more potential phage particle profiles, especially in ecology or antibiotic therapy involving reasonable usage. Four main aspects are currently constituting the main framework of phage research: phage identification, species classification, lifestyle, and host relationship. Deep learning frameworks (convolutional neural networks, graph neural networks, Transformer, etc) are often used to assist in those aspects, which provide potential biological insights. Five cases are provided in this section (Figure 4), first, the method of using protein clusters as tokens in Transformer for phage identification is recomposed in MicroWorldOmics [38], aiding the identification of phage sequences in metagenomic data. Then, using a graph convolutional network for phage taxonomy assignment [39], combining self-supervised/fine-tuning to distinguish phage lifestyle [40], and using a knowledge graph to assist phage host prediction [41] are all included in MicroWorldOmics as a separate plugin. In order to overcome the problem of time, resource consumption, and platform incompatibility in deep learning method, some software (diamond, blast) [25, 42] was changed to Windows version, and some algorithms such as Markov cluster algorithm (MCL) was rewritten to be platform compatible and reduce computation. Compared with Linux (with 16-core CPU and 62GB memory), plugins “Cherry” and “Phagcn” performed less time consumption (Phagcn average savings: 500 s, Cherry average savings: 1700 s) in Windows (with 2-core CPU and 16GB memory) (Figure S5 A and B). Because of platform compatibility, the recompiled plugins “Phamer” and “Phatyp” will increase the extra running time (Phamer average increases: 140 s, Phatyp average increases: 100 s) (Figure S5 C and D). For the plugin “Genomad”, software to identify mobile elements [43], which cannot be compiled in the window system and carries its own database, it will be deployed in a remote server. Users only need to submit files through the MicroWorldOmics interface and wait for the result to be sent from the remote server. Roughly the same time is spent between used in Windows and used in Linux (Figure S5 E).

**Figure. 4.**
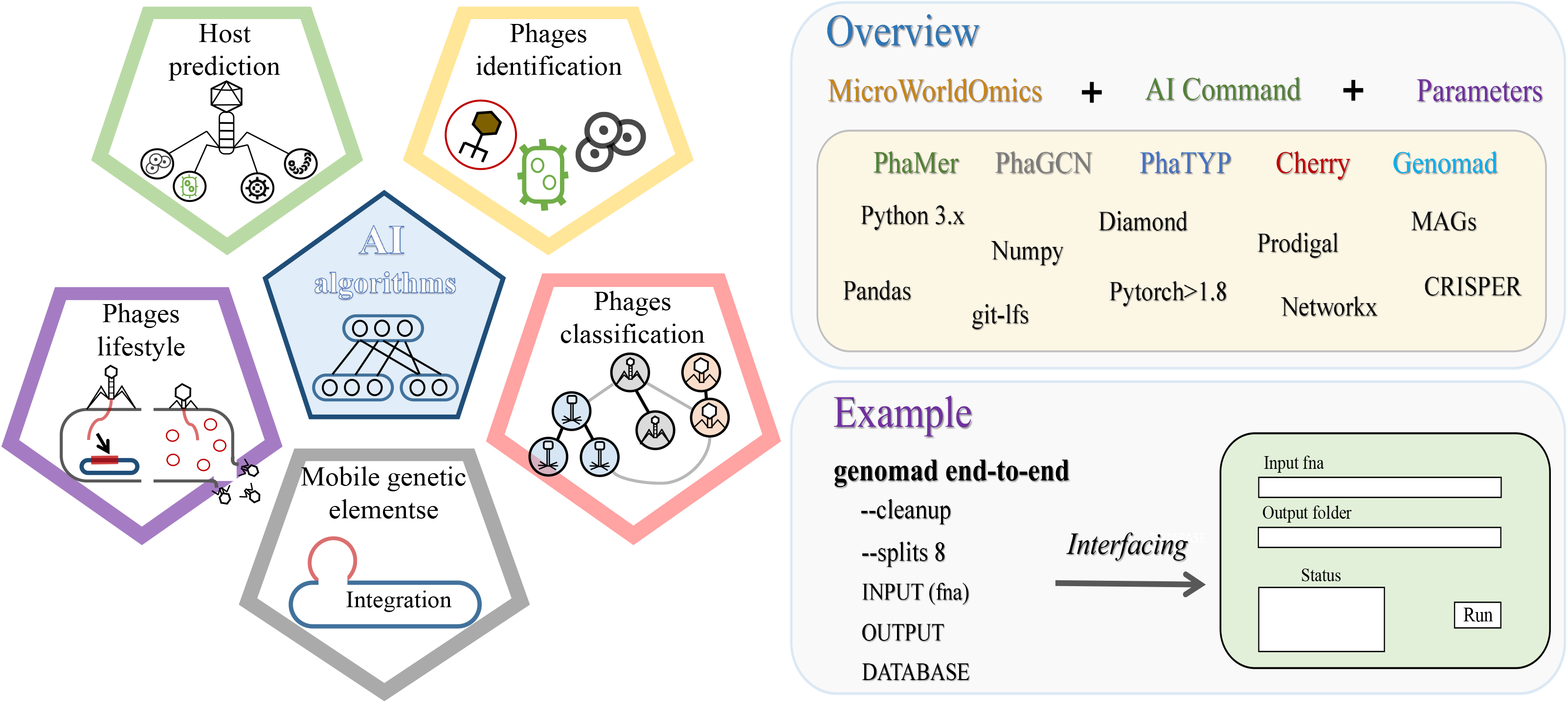
Using deep learning models to assist virome data analysis, including identifying phage sequences, phage taxonomy assignment, phage life style identification, phage host prediction, and search for mobile elements. All analysis methods and deep learning frameworks are provided in user-friendly plugins.

### Case 3: Isolates genome-based and metagenome-based data analysis and visualization

In the MicroWorldOmics framework, the analysis of bacteria is divided into two modules: 1. analysis of isolates based on whole genome sequencing; 2. Microbial flora analysis based on metagenomic or amplicon sequencing. The advantage of whole genome sequencing is that isolates can be obtained with complete sequence information, geographical distribution, evolutionary relationships between isolates, tracing inferences, and identification of gene islands are the main focus of the analysis process. Based on the global map information that has been collected, the geographic distribution of the sampled strains can be displayed. The “ShinyMap” plugin saves the geographic information covering major countries in the world into RData form (stored in the demo), and users only need to edit the metadata collected by the corresponding region and upload it to complete the visualization. Of note, the depth of color represents the number of samples collected (Figure 5A). Core genome in isolates, the more conservative elements under the same genus, which is the high-frequency concept to demonstrate evolutionary relationships (see Materials and methods). On the other hand, if the core fragment is judged too strictly, the parameters “Identification” and “Gene coverage” can be freely defined by users (Figure 5B). Besides, we embed “BactDating” [44] into MicroWorldOmics by Shiny, which is based on the R language inference molecular clock model. Based on the Bayesian algorithm, this plugin provided an interface to obtain temporary signal and time tree through functions “roottotip()” and “bactdate()” (Figure 5C and D). Furthermore, this application based on the Bayesian algorithm also can be performed to analyze population structure (use parameters “max depth” and “max number of populations” to control the details of the cluster). In actual projects, branch-ignoring and branch-considering trees are provided in this plugin (Figure 5E and F).

**Figure. 5.**
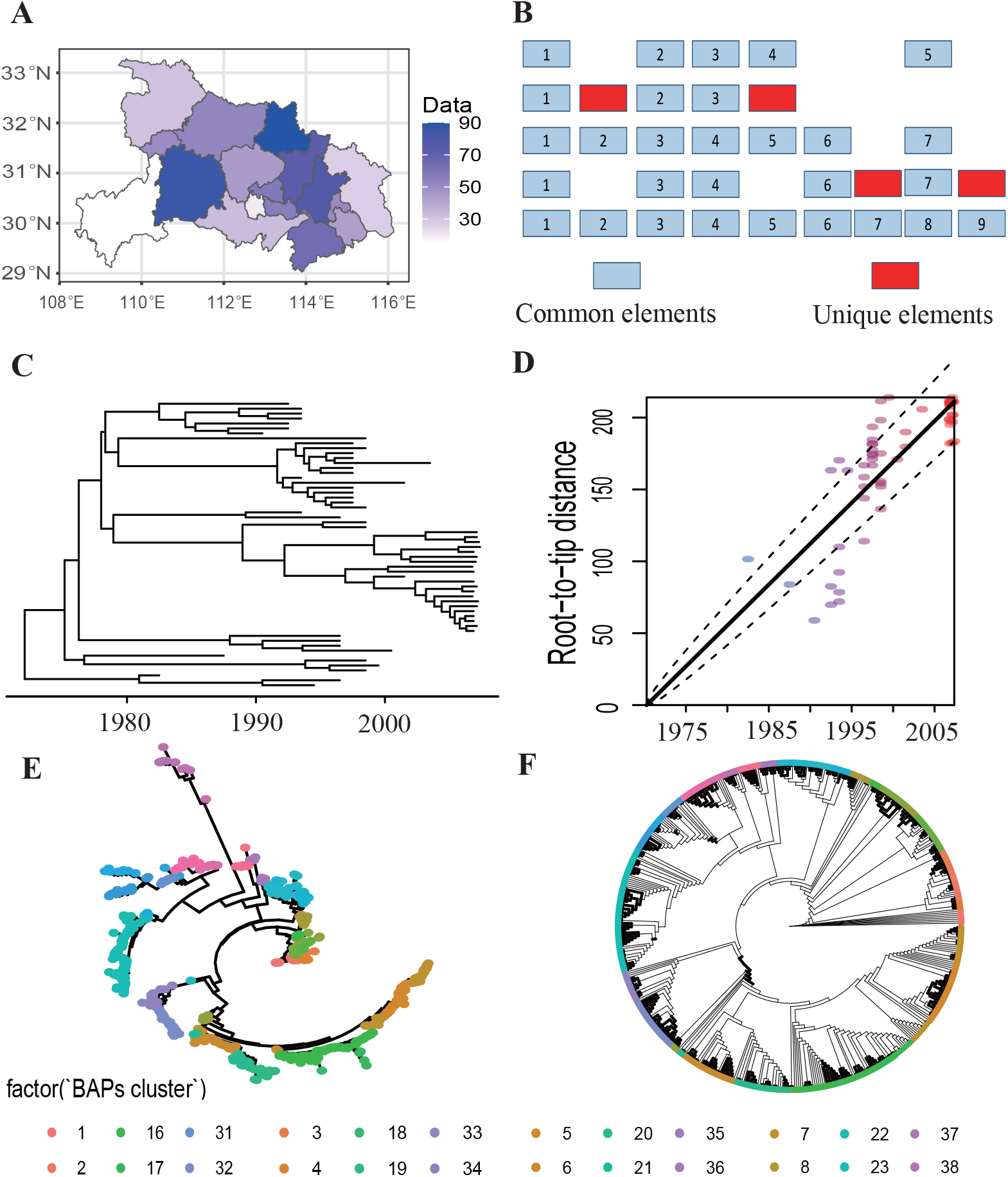
Representative use cases of whole isolates genome analysis in MicroWorldOmics. (A) Maps can be used to display sampled information. (B) In the construction idea of the core genome, the red elements represent the unique elements in isolates, and the blue elements represent the shared elements of the isolates. (C) The plugin “ShinyBactDating” is used to calculate the molecular clock model of isolates for inferring the differentiation time. (D) The temporary signal is calculated using the function “roottotip()” of the plugin “ShinyBactDating” (rate = 10.4, MRCA = 1978.1, R^2^ = 0.97, p < 1e-4). Bayes method can be based on the needs of users for different depths of clustering, support branch-considering (E) and branch-ignoring (F) trees.

Most studies utilizing “microbiome” or metagenomics techniques to date have considered the discovery and identification of new microbial populations, discussion in population relationships, and the regulation between population and ecological environment or health. Dozens of metagenomics/amplicon analysis plugins have been developed in MicroWorldOmics. In 2023, Tao Wen et al. summarized the best practice for microbiome analysis by R [45] and released the source code. This study implemented the Shiny-based web interface based on released the source code, allowing users to practice, download, explore, and quickly analyze/visualize the data set from Tao Wen et al. Here, we used the Metagenomics/Amplicons module of MicroWorldOmics to show that the excellent data analysis and visualization were significantly convenient for researchers without programming. Among the dimensionality reduction results, the PCoA plot shows the clustering of samples by distance matrix (Figure 6A). The difference between groups is higher than the inter-individual variation, that is, while microbial communities between different groups were scattered, the same group (in KO, OE, or WT) was closely gathered except for several samples. In the same way, the microbial abundance matrix can be presented after Non-metric multidimensional scaling (NMDS) in each sample (Figure 6B). Ternary plots, describe the proportional relationship between three variables for microbial abundance, the plugin is allowed to select any microbial classification level (kingdom, phylum, class, order, family, genus, and species) to show the proportion of microbial flora in the three groups (Figure D). Benefiting from the powerful R package <u>ggClusterNet</u> [46] and Shiny convenience operation for network visualization, based on the functional interface provided by <u>ggClusterNet</u>, MicroWorldOmics supports microbial network visualization (Figure C and E). The level of microbial classification (kingdom, phylum, class, order, family, genus, and species) can be arbitrarily selected by the user through the dropdown option box on the Shiny panel. In order to ensure the beautiful and publish ability of the network diagram, the plug-in also provides a custom color system (R package ColourPicker) for the beautification of the figures. Besides, some advanced analytics plugins are also included in MicroWorldOmics, such as the time series analysis plugin “ShinyTimeSeries”, plugin “Bugbase” for prediction of bacterial phenotype based on 16S OTU table, plugin “ShinyBatch” to remove the microbial batch effect, etc (see Table S3).

**Figure. 6.**
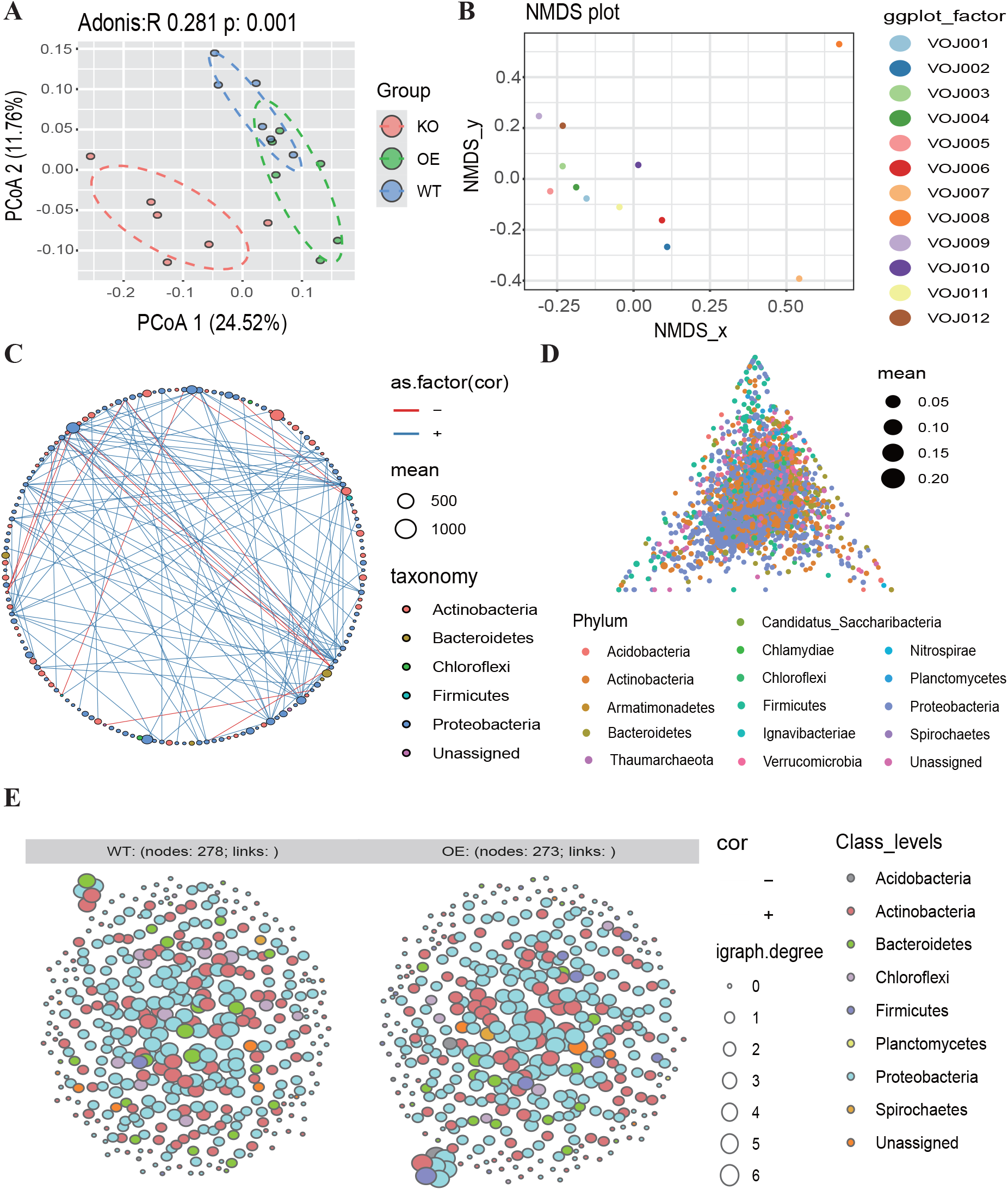
Representative use cases of metagenomic/amplicons analysis in MicroWorldOmics. (A) Based on the microbial abundance matrix, three groups in the PCoA dimensionality reduction cluster showed good differentiation (PCoA 24.52%, PCoA 11.76%). (B) NMDS can be analyzed between samples. (C) Microbial interaction networks constructed by using correlations (Pearson or Spearman correlation coefficient) can be displayed in different microbial category levels. The category level can be selected in the Shiny panel by dropping down boxes. (D) The ternary phase diagram can show the proportional relationship of different microbial content in the three groups, and obtain the dominant microbial flora. (E) Visualize microbial networks with custom colors and groups.

### Case 4: Network comprehensive construction

Microbial communities have complex community relationships, including saprophytic, parasitic, symbiotic, and other diverse survival strategies. Graphs, a data structure that represents node relationships, play an important role in measuring microbial community interactions. Based on the construction method, they are divided into four categories: 1. Correlation & regression-based; 2. Graphical model inference; 3. Cosine- based network; 4. Differential network analysis (Figure 7A). Those methods can be implemented in advanced plugins, such as ShinyMicroWGCNA, ShinySpiecEasi, ShinyREBACCA, ShinyCCLasso, ShinyCosine, and ShinyDiffCoEx, to construct different networks and display their weights between nodes. In addition to constructing interaction networks, network characteristics are also used interactively to search for microbial markers under different conditions, for example, R package NetMoss2 [47] provides a comprehensive analysis and visualization function based on the network structure of microbiota, then on this basis, interactive applications are provided by Shiny. Obviously, the top 30 microflora with the highest NetMoss score can be clearly identified, hence, Lachnospira is the highest one with significant enrichment in control (Figure 7B). Analysis of traits associated with the microbiome has also been applied in MicorWorldOmics as a complete pipeline based on the basic principle of the WGCNA package [33], including soft threshold filtering, One-step network construction, modules and traits relationship, export modules, and network heatmap plot those 5 steps. Eventually, ME_4 is highly correlated with D6 (corr = 0.65, pval=5e-04), indicating that the microflora of this module may affect the occurrence of trait D6 (Figure 7C). Compared to identify differences in individual microflora, especially in the study of the gut microbiome, has increasingly failed to meet the needs of scientific research. When researchers are faced with a complex microworld system, the perspective of overall regulation needs to be proposed to solve biological problems. Differential networks are often used to mine differences in the microbiota regulatory patterns under two conditions. Based on this method (see Materials and methods), modules with different regulatory patterns can be clearly found under the two conditions, the regulatory relationship in M1, M2, and M5 is more sparser in condition A than in condition B. On the contrary, the regulatory relationship in M4 is more closer in condition A (Figure 7D).

**Figure. 7.**
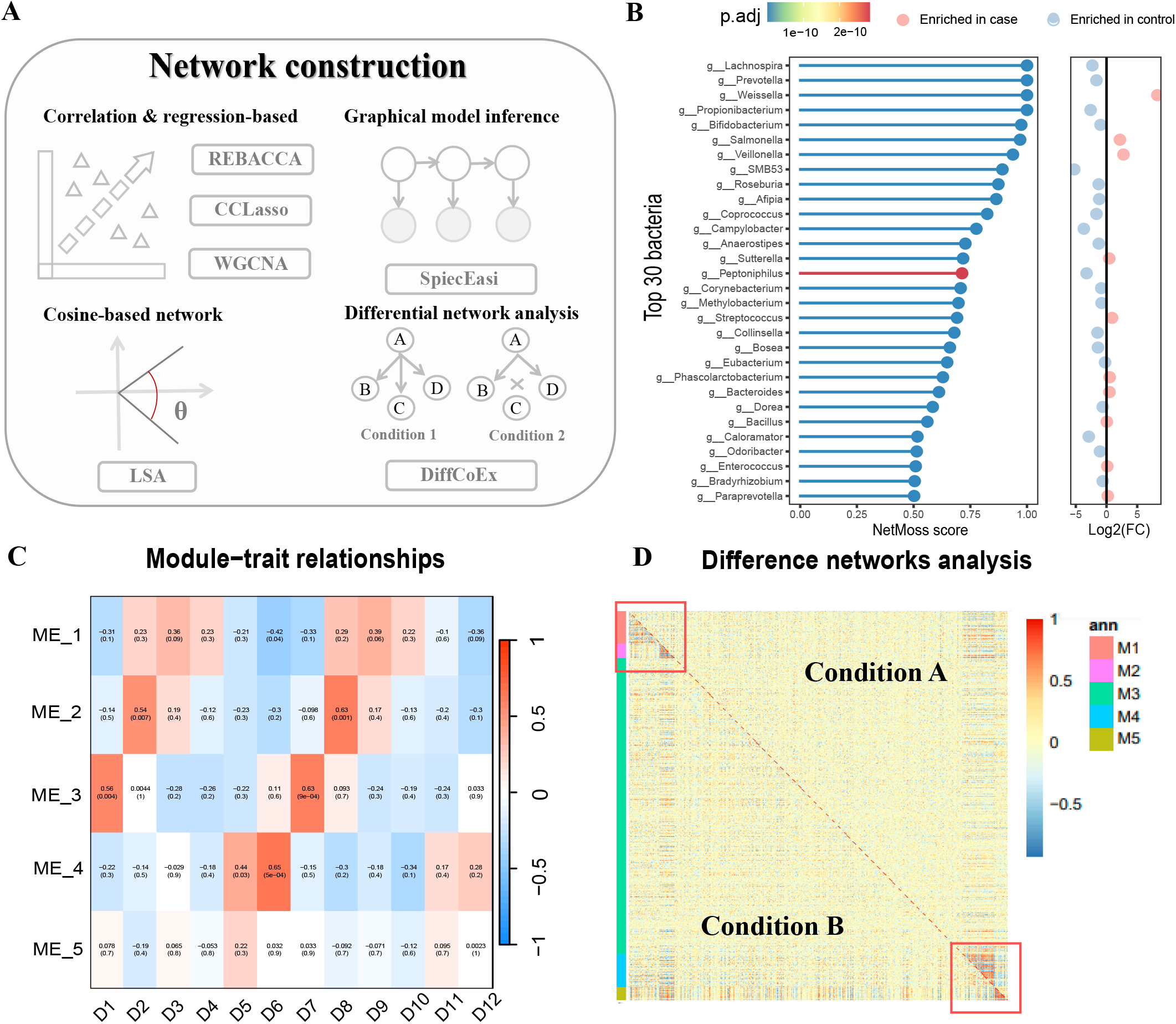
Representative use cases of network analysis in MicroWorldOmics. (A) There are four types of network analysis that are offered to users, including correlation & regression-based method, graphical model inference, cosine-based network, and differential network analysis. (B) The microbial network is used to find the marker microbial flora. The figure shows the top 30 microbial flora with the highest NetMoss scores (Lachnospira is the highest). The panel on the right provides the enrichment degree of the flora in the comparison group. (C) Based on the basic principle of WGCNA, the correlation analysis between microbial flora and trait was carried out. As shown in the figure, the microbial flora of the M4 module had the highest correlation with trait D6 (corr = 0.65, p_val = 5e-04). (D) Differential network analysis, looking for the interaction network of microbial flora differences in conditions A and B, as shown in the figure, M1, M2, and M5 in the red box have a stronger regulatory relationship in condition B, while M4 has a stronger regulatory relationship in condition A.

## Discussion

In summary, facing for microbiome and virome comprehensive analysis methods are integrated into MicroWorldOmics, containing 90 sub-applications in 14 modules (Seven of them are self-developed, and the rest are included in public software). More than 80 Python modules and 600 R packages are invoked, scoping genome sequence processing, alignment, deep learning, mathematical statistics, dimensionality reduction, difference analysis, batch effect, normalization, visualization, etc. A good interactive development experience is given to developers by PYQT5 and Biopython and Shiny framework also creates smart components for user interface. Meanwhile, multiple data input and output formats are supported, such as fasta, GFF, TXT, CSV, PNG, JPG, etc. MicroWorldOmics focuses on providing a convenient, one-stop analysis process, the emergence of this model has greatly saved the time of researchers, allowing researchers to focus more on the study of microbes and health, virus and host co-evolution, and another related issue.

MicroWorldOmics provides a download channel on Github (Github:https://github.com/hzaurzli/MicroWorldOmics) and an opening Community Forum, a user-facing communication channel to better maintain MicroWorldOmics.

From another perspective, the rapid development of sequencing technology has led to drive the development of bioinformatics software. At present, a large number of bioinformatics software requires users to have a certain programming foundation. Nevertheless, the high-frequency demands of scientific research users, convenient scientific analysis pipeline, and more interactive data mining applications are considered of great importance in promoting data science for biomedical researchers. Here, some excellent software has been developed to eliminate the gap caused by tedious computer knowledge, such as TOmicsVis for RNA-seq analysis, Hiplot for lightweight visualization, PhyloSuite is responsible for evolutionary analysis, and TBtools, OmicsSuite are used for plant and horticultural studies. The emergence of MicroWorldOmics fills the gap of microbiome and virome data mining in this kind of research and MicroWorldOmics has minimized data analysis tasks’ learning and usage costs for users without programming skills. However, there are still some contradictions between supply and demand in the microbiome data analysis, difficulty in controlling the space size of the deep learning plugin under the Windows system, some plugins running slowly, and a large number of upstream analysis tasks lacking in MicroWorldOmics still need to further development. Although the various problems mentioned above, existing analysis plugins deployed on this software have greatly enhanced the operability and maintainability. Moreover, more analysis methods or frameworks (In particular, the analysis of correlations between microbes and traits, or new artificial intelligence frameworks) will be implemented in the future. Then, these resources will be fully utilized in MicroWorldOmics, bringing users a new experience.

## Codes availability

The software resource codes are available on GitHub: https://github.com/hzaurzli/MicroWorldOmics.

## Contributor Information

Runze Li, National key laboratory of agricultural microbiology, College of Biomedicine and Health, Huazhong Agricultural University, No. 1, Shizishan Street,Wuhan, Huazhong Agricultural University, Wuhan, 430070, China.

## Acknowledgment

This research was supported by the National Natural Science Foundation of China (32072323, 32073022, 31772083), the National Key Research and Development Program of China (2022YFD1800900), Training Program of Distinguished Agricultural Researcher supported by the Ministry of Agriculture and Rural Affairs (13210333), HZAU-AGIS Cooperation Fund (SZYJY2022018, SZYJY2022027), the National Innovation and Entrepreneurship Training Program for Undergraduates (S202210504224, 2022310, S202010504215, 202110504076), the Natural Science Foundation of Hubei Province (2022CFB659, 2023AFA111) and The Young Top-notch Talent Cultivation Program of Hubei Province.

## Supporting information

The online version contains supplementary figures and tables available.

---

**Figure. S1.**
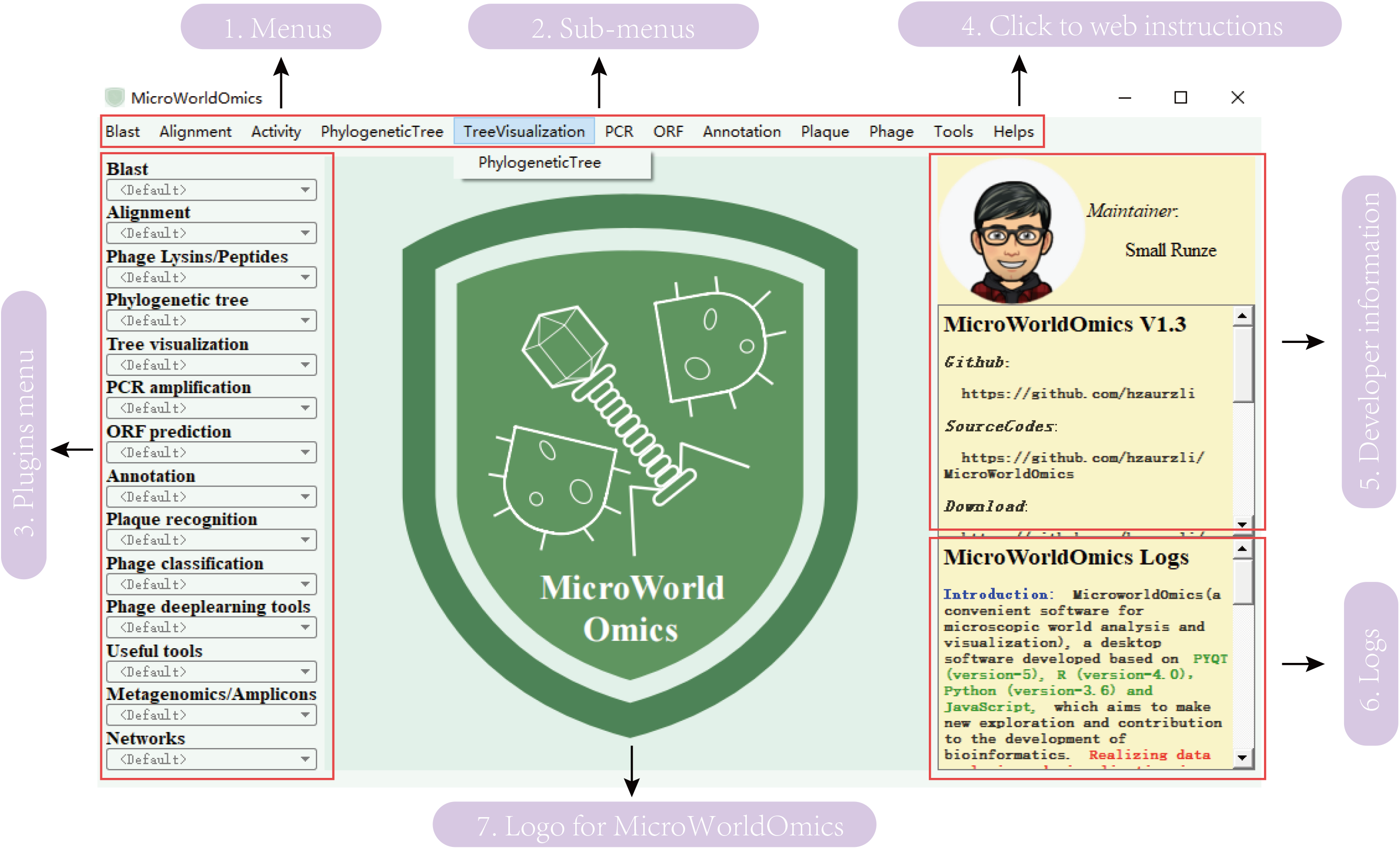
MicroWorldOmics layout and sub-application user interface. The layout of MicroWorldOmic’s main page and the sub-application user interface are distributed in the top and left columns. The middle represents the logo, and the right column represents the developer and the development log.

**Figure. S2.**
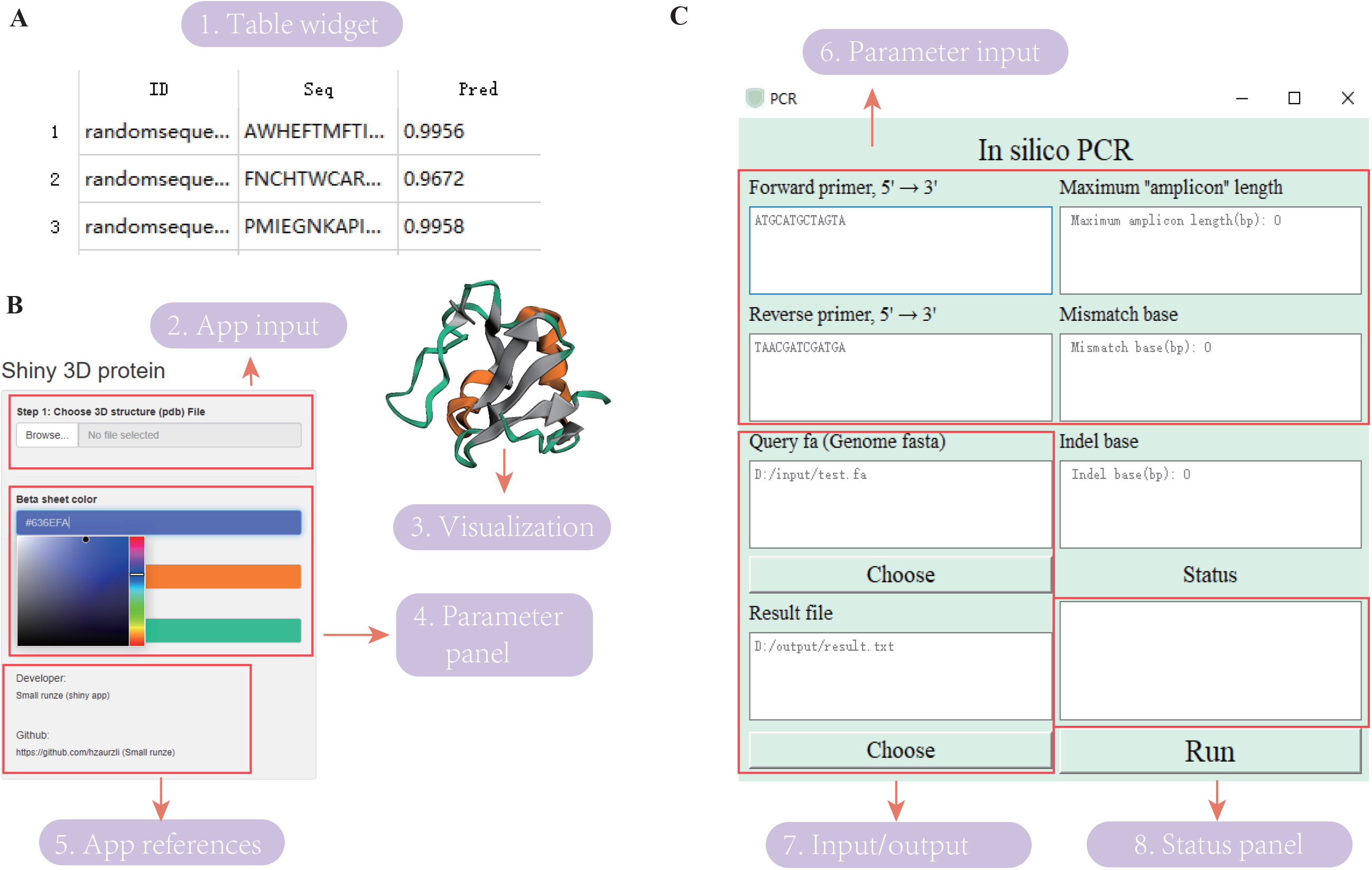
Plugin function page, including table display (A), Shiny app usage details and parameter settings (B), widget panel display, and parameter settings (C).

**Figure. S3.**
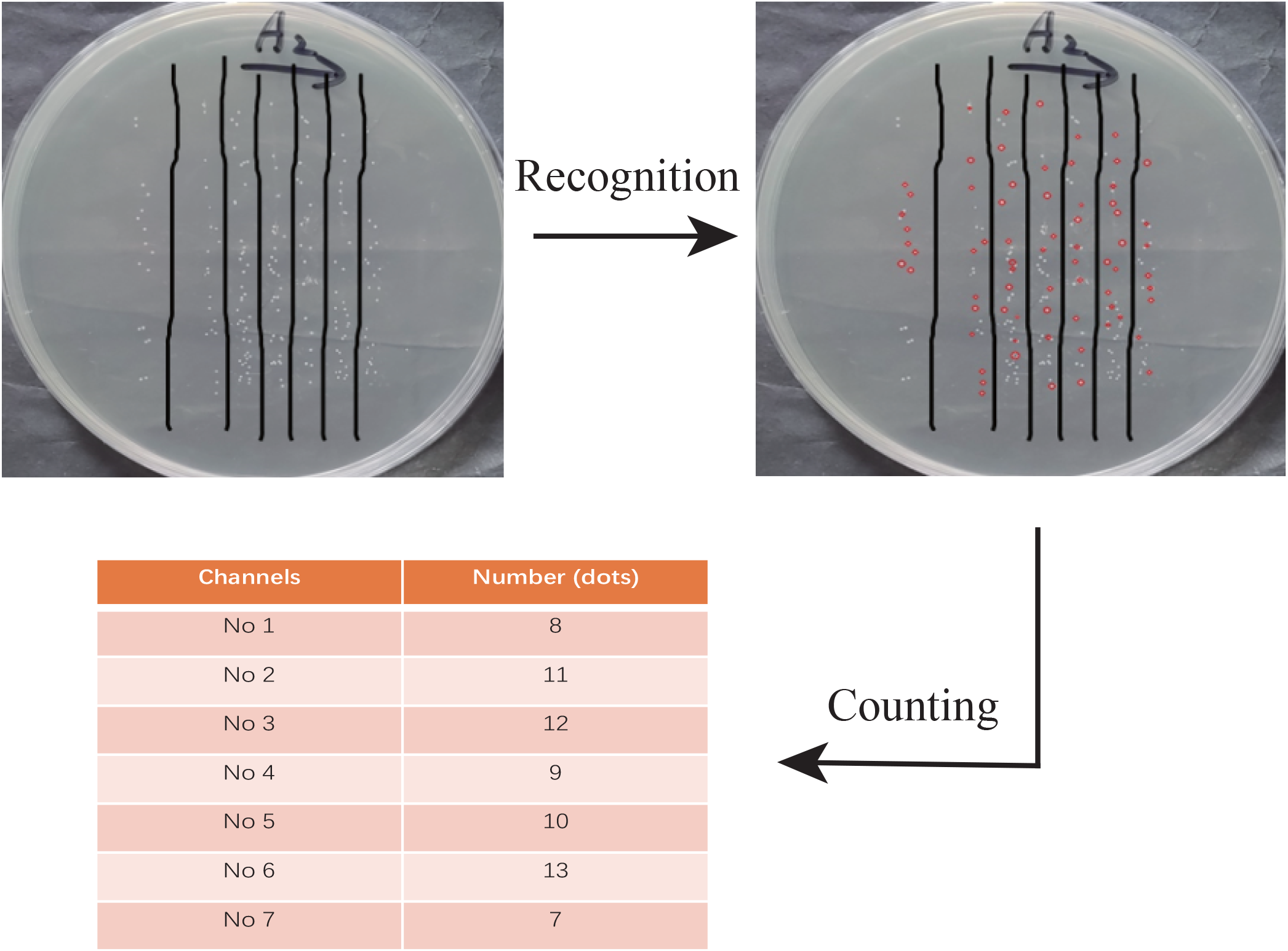
Strain identification function can be more convenient for plaque identification, and can also be divided into channels to identify plaque.

**Figure. S4.**
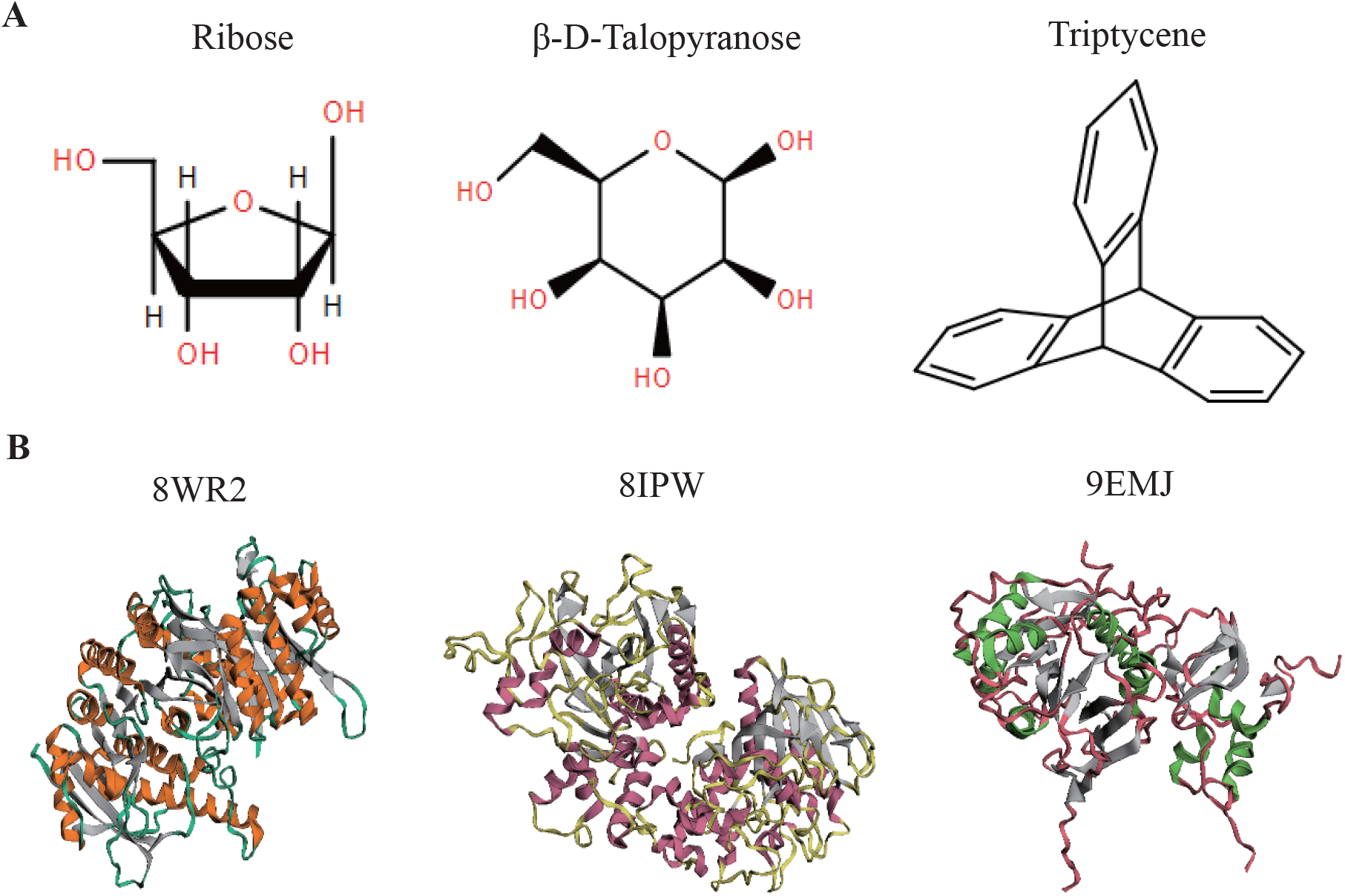
Visualization function of protein three-dimensional structure (A) and chemical formula structure (B).

**Figure. S5.**
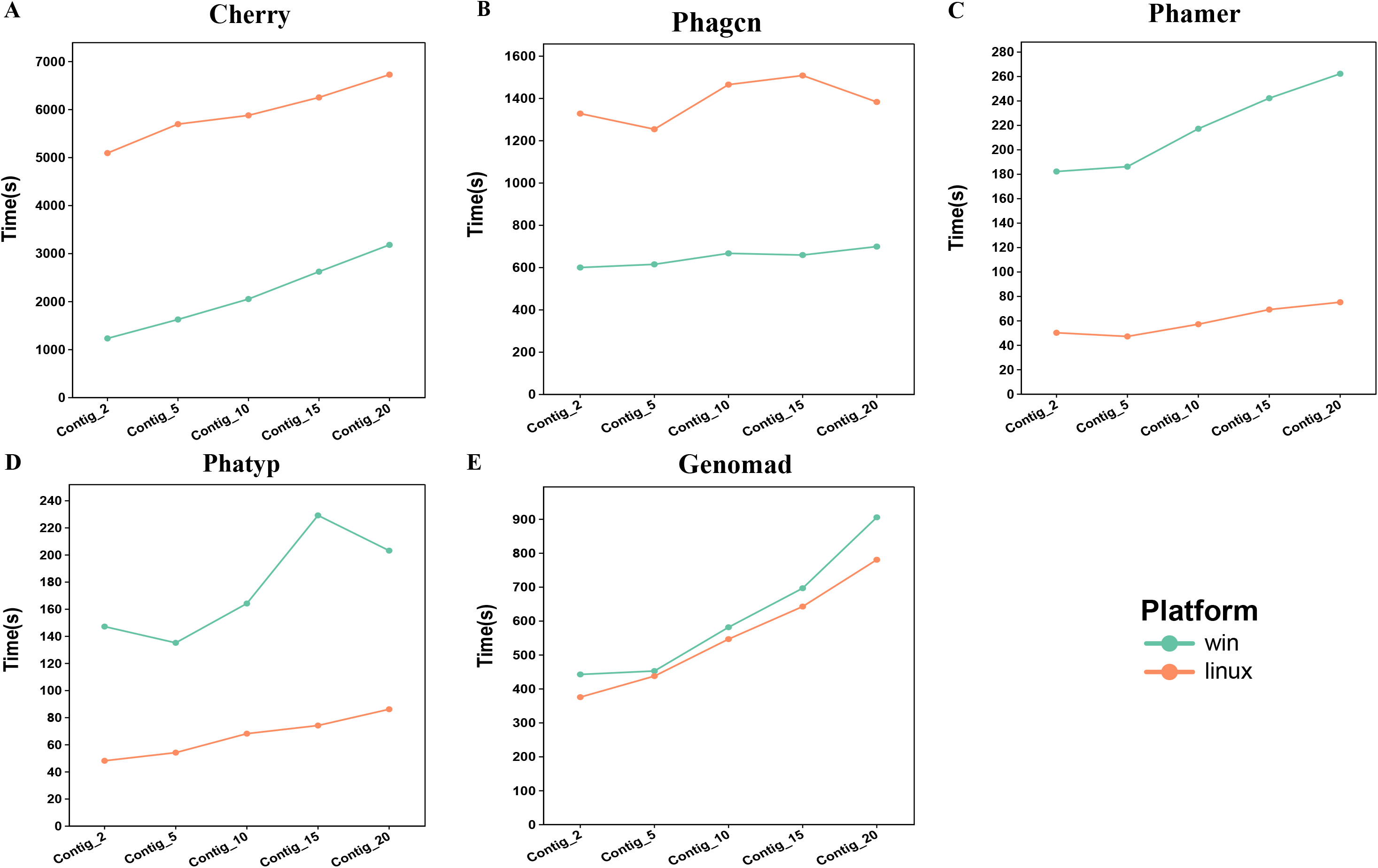
Time consumption of deep learning models in assisted virome analysis. In phage taxonomy assignment (Phagcn) and phage host prediction (Cherry), the runtime on Linux (with 16-core CPU and 62GB memory) is greater than on Windows (with 2-core CPU and 16GB memory) (A, B). While, in identifying phage sequences (Phamer) and phage lifestyle identification (Phatyp), the runtime on Windows (with 2-core CPU and 16GB memory) is greater than on Linux (with 16-core CPU and 62GB memory) (C, D). Finally, the plugin “Genomad” runs roughly the same time on Linux and Windows (E).

**Table S1.** Basic models used in lysin activity evaluation.

| Model | Param |
| --- | --- |
| Extremely randomized trees | ERT |
| Logistic regression | LR |
| Artificial neural network | ANN |
| Extreme gradient boosting | XGB |
| K-nearest neighbor | KNN |

**Table S2.** Feature types used in lysin activity evaluation.

| Feature | Param |
| --- | --- |
| Amino acid index | AAI |
| In grouped amino acid composition | GAAC |
| Composition–transition–distribution | CTD |
| Amino acid entropy | AAE |
| Amino acid composition | AAC |
| Binary Profile-basedfeature | BPNC |

**Table S3.** Metagenomics/Amplicons module functions.

| Plugins | Functions |
| --- | --- |
| ShinyMultifun | Comprehensive analysis module<br>(diversity analysis, dimensionality reduction,<br>normalization) |
| ShinyDiff | Differential analysis<br>(Maaslin2 [48], based on mixed linear model) |
| ShinyNMDS | Non-metric multidimensional scaling<br>(NMDS) |
| ShinyPCoA | Principal co-ordinates analysis<br>(PCoA) |
| ShinyMCA | Multiple correspondence analysis<br>(MCA [49]) |
| ShinyVolcano | Visualization of volcano map |
| ShinyTimeSeries | Microbial abundance time series analysis |
| ShinyBatch | Remove microbial data batch effect |
| Bugbase | Prediction of organism-level microbiome<br>phenotypes [50] |
| ShinyNetMoss | Biomarker identification through microbial<br>networks [47] |

